# *K*-mer similarity, networks of microbial genomes and taxonomic rank

**DOI:** 10.1101/125237

**Authors:** Guillaume Bernard, Paul Greenfield, Mark A. Ragan, Cheong Xin Chan

## Abstract

Alignment-free (AF) methods have recently been adopted to infer phylogenetic trees. However, the evolutionary relationships among microbes, impacted by common phenomena such as lateral genetic transfer and rearrangement, cannot be adequately captured in a strictly tree-like structure. Bacterial and archaeal genomes consist of highly conserved regions, e.g. ribosomal RNA genes (commonly used as phylogenetic markers), more-variable regions and extrachromosomal elements, i.e. plasmids (that contain genes critical under a selective condition e.g. antibiotic resistance). The impact of these elements on genome-scale inference of microbial phylogeny remains little known. Here, using an AF approach, we inferred phylogenomic networks of microbial life based on 2785 completely sequenced bacterial and archaeal genomes, and systematically assessed the impact of ribosomal RNA genes and plasmid sequences in this network. Our results indicate that k-mer similarity can correlate with taxonomic rank of microbes. Using a relational database approach, we linked the implicated *k*-mers to annotated genomic regions (thus functions), and defined core functions in specific phyletic groups and genera. We found that, in most phyla, highly conserved functions are often related to Amino acid metabolism and transport, and Energy production and conversion. Our findings indicate that AF phylogenomics can be used to infer reticulate relationships in a scalable manner and provide new perspective into microbial biology and evolution.

## Introduction

Genome evolution in microbes involves highly dynamic molecular mechanisms including genome rearrangement and lateral genetic transfer (LGT). These mechanisms may violate the implicit assumption of full-length contiguity in multiple sequence alignment (MSA), a common step in phylogenetic analysis. Furthermore, MSA-based approaches necessitate heuristic methods e.g. Bayesian inference in reconstructing phylogenies, which are not scalable to the quantity of existing and forthcoming genome data^1,2^. An alternative strategy is to infer evolutionary relatedness based on shared subsequences of fixed length, known as *k*-mers, i.e. *alignment-free* (AF) methods^3^. AF approaches provide exact solutions (i.e. pairwise distances between genomes based on shared *k*-mers) which can be directly used in deriving a phylogenomic network^4^.

In the past decades, AF approaches have been used in phylogenomics to infer phylogenetic trees of evolving sequences^2^, complete genomes^5-7^ and NGS data^8^. The AF approaches used in phylogenomics can be classified into two categories^3^, one based on the count of *k*-mers^9,10^ and the other based on match lengths^11,12^. Methods in both categories have been shown to be scalable and accurate in inferring phylogenies at both gene and genome level^2,10^ while being robust to complex evolutionary event such as LGT or rearrangement^7^.

However, the evolution of microbial genomes is known to not follow a tree-like structure, notably because of widely spread LGT events^13,14^, and a network structure to represent phylogenies has been increasingly chosen over a traditional tree representation since the emergence of genome data^15^. Different types of phylogenetic networks have been developed to increase our understanding of microbial evolution, including genome networks^15^ and sequence-similarity networks^16^. These networks, by allowing more than one connection per node, i.e. sequences or genomes, can be used to visualise vertical phylogenetic signal, from an inferred (but un-observed) common ancestor to organisms, and lateral signal between organisms. But the phylogenetic signal used to build these networks is generally inferred using BLAST hits^17^ and, therefore, based on sequence alignment.

As a proof of concept, we previously generated an AF phylogenetic network for 143 bacterial and archaeal genomes^18^, using pairwise distances based on the 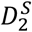 statistic. By varying similarity thresholds in displaying the network, we could easily capture changes in the network structure, e.g. cliques, which reflect evolutionary events and dynamics of microbial genomes. For instance, we recovered the progressive separation of the different genomic lineages throughout their evolution^18^ and showcased particular relationships between isolates not observed using a classical tree structure^7^.

Highly conserved regions such as rRNA genes have long been used as phylogenetic markers for the inference of trees, and indeed our current view of the Tree of Life is based on ribosomal proteins^19^. However, trees based solely on this or other markers represent only a small fraction of the total genomic information^20^. On the other hand, variable regions or exogenous genetic material, i.e. plasmid genomes, are rarely taken into account when inferring phylogenies. Plasmids genomes are known to be important agents of LGT in microbes^21^,^22^ as well as a common vector of antibiotic resistance^23^, and a better understanding of their contribution to microbial evolution is an urgent matter due to the widespread of antibiotic resistance^24^. The AF approaches allows for the comparison of whole genomes, including exogenous sequences, with good computational performance but they do not keep information related to the *k*-mers locations^3^,^7^. Without positional information, the contribution of specific regions, such as rRNA genes or those having arisen from exogenous genetic material, to the phylogenetic signal captured by AF methods remains difficult to assess.

Here, to investigate the impact of plasmids and highly conserved genes in phylogenomic inference, using 2785 complete bacterial genomes we inferred AF phylogenomic networks using (a) all genome data including plasmids, (b) chromosomal sequences without ribosomal RNA genes, (c) only ribosomal RNA genes and (d) only plasmid sequences. We systematically assessed the impact of rRNA genes and plasmids on the overall microbial phylogenomic network. Using an advanced database approach, we investigated the core functions that are specific to particular phyletic groups or genera based on the shared *k*-mers.

## Results

To infer a phylogenomic network, we first calculated a pairwise distance *d* between each genome based on the 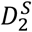 distance using 25-mers (see Methods). For each pair of genomes *a* and *b* we transformed *d_ab_* into a similarity value *S_ab_* and generated a similarity network, following our earlier approach^18^; we consider this network to depict phylogenetic relatedness among these genomes, i.e. to be a phylogenomic network. Here we define a threshold *t* for which only edges with *S* ≥ *t* are considered in the network. To compare our results at the genome and phylum levels, we generated *I-*networks in which a node represents a distinct genome isolate and an edge between two nodes (isolates) indicates evidence of shared *k*-mers, and *P*-networks in which a node represents a distinct phylum and an edge represents the number of isolates that share *k*-mers with isolates of another phylum (see Methods). We then compared the k-mer networks based on the topological differences between them at different *t*. All the *I*- and *P*-networks of these 2705 genome isolates are available at http://espace.library.uq.edu.au/view/UQ:543037^25^.

### AF networks of microbial evolution

To infer a phylogenomic network of prokaryotes, we used a dataset of 2785 completely sequenced microbial genomes (2619 Bacteria, 176 Archaea) as of 31 January 2016 (Supplementary Table S1). To eliminate redundancy among the data, we kept only one genome where an identical genome (from another isolate) was present (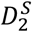 distance = 0). We also removed genomes with little evidence of shared *k*-mers (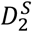 distance > 10); these genomes share ≤ 0.01% of 25-mers with any other genomes (i.e. there is little evidence of homology). Following this filtering step, we took a total of 2705 genomes forward into subsequent analyses. For each network, we systematically assessed the number of connected nodes ( *c*), number of edges ( *e*), maximum clique size ( *z*) and maximum number of cliques ( *n*) across varying levels of the similarity-score threshold *t*. Here we required a clique to contain three or more edges, and we defined *E* as the average number of edges per node.

The network topology changes substantially with similarity threshold: at *t* = 0, *c* = 2705, *e* = 3835070 and *z* = 2700, compared to *c* = 1358, *e* = 9898 and *z* = 48 at *t* = 9 (Supplementary Table S2). As we required more-stringent threshold of shared similarity, the network became less-connected, and distinct cliques corresponding to diverse taxa (i.e. phyla, classes, genera) started to form. For example, Bacteria and Archaea form distinct cliques at *t* = 4, most phyla can be identified as distinct cliques or paracliques at *t* = 5, and all proteobacterial classes are separate from each other at *t* > 5.

The *I*-network is very densely connected at *t* = 0, with the maximum number of cliques *n* = 10. The value *n* is too high to be computed at *t* = 1 or *t* = 2, but is 1662785 at *t* = 3 and decreases to 232 at *t* = 9 (Supplementary Table S2). Most isolates are members of a single large clique at *t* = 0 and *t* = 1, in which *E* > 1400; at *t* = 2, *E* = 736.3. The network becomes less dense at *t* = 3, with *E* = 112.8 (Supplementary Table S2). As this network of 2705 nodes remains too densely connected to be visualised and analysed directly, we generated the P-network using the same data, with each node representing a phylum. Figure 1 shows the *P*-network of the 2705 genomes at *t* = 3. The thickness of each edge represents the number of instances in which any two genomes (one for each phyla connected by the edge) have a similarity value *S* ≥ *t*. Major phyla (e.g. β- and γ-Proteobacteria, Firmicutes, Actinobacteria and Tenericutes) are clearly separated at *t* = 3. The thickest edge (in yellow) is between the β-Proteobacteria and *γ*-Proteobacteria, suggesting a high similarity among genomes between these groups. In addition, we also observed a large extent of shared 25-mers between Firmicutes and any of the proteobacterial classes.

**Figure 1:**
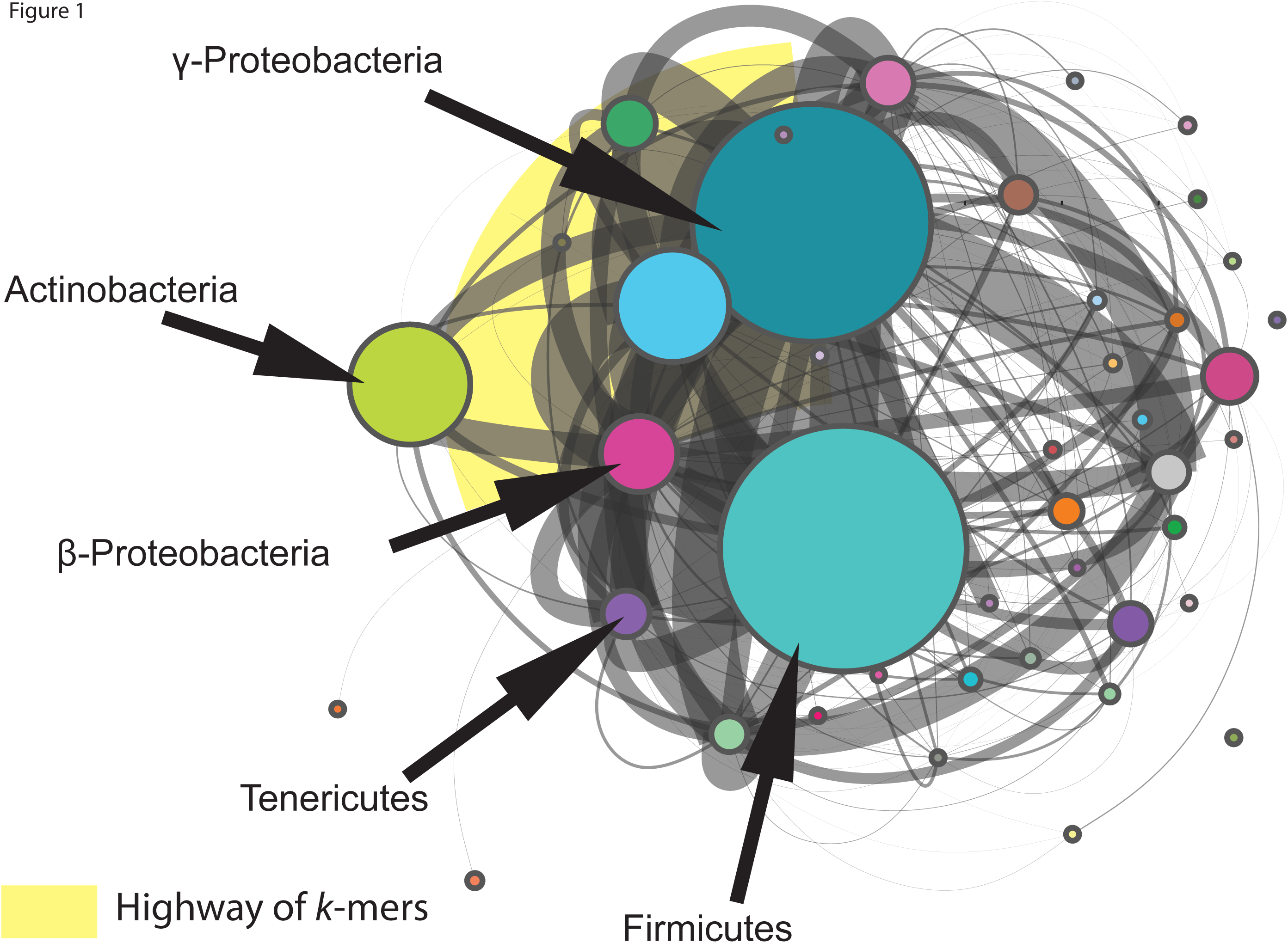
*P*-network of prokaryote genomes using 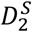 with *k* = 25 based on whole-genome data, at *t* = 3. An edge between two nodes represents the number of connections between isolates from the two phyla. The size of a node is proportional to the number of isolates within the phylum.

### Impact of rRNA genes

To determine the contribution of the highly conserved rRNA genes to our AF networks, we computed AF networks using 2616 genomes (a subset of the 2705 above) upon excluding all rRNA gene sequences, and genomes with no annotated information (see Methods). The *I*-network of the genomes from which rRNA genes have been removed has a lower density than the one inferred using the whole dataset. Similarly to the previous *I*-network, here at *t* = 0, *c* = 2615, *e* = 1720082 and *z* = 1226, and these values decreased to *c* = 1290, *e* = 9008 and z = 47 at *t* = 9 (Supplementary Table S3). At *t* = 3, the *I*-network of the rRNA gene-free network has 38.9 edges per node on average, about 2.9-fold fewer than the 112.8 edges per node in the whole-genome network (Supplementary Table S2). Figure 2 shows the *P*-network of these 2616 genomes at *t* = 3. As in Figure 1, the thickest edge (in yellow), between β- and γ-Proteobacteria (Figure 2), indicates the largest number of instances of shared *k*-mers between genomes from these two groups. This *P*-network is less dense than the equivalent network based on the whole data (shown in Figure 1). Although we observed fewer connections between the phyla after removal of rRNA sequences from the genome data, many of the major connections observed in Figure 1 remain, *e.g*. between β- and γ-Proteobacteria, and between Actinobacteria and *γ*-Proteobacteria. Thus the sharing of 25-mers contributing to these major connections extends beyond the commonly used phylogenetic marker of rRNA genes.

A network computed using only the rRNA genes sequences (see Methods) was denser than the two corresponding *I*-networks above. At *t* = 6, *E* is high at 854.4 ( *z* = 1321; Supplementary Table S4), compared to 10.4 ( *z* = 82) and 9.6 ( *z* = 74) in the *I*-networks based on whole-genome and rRNA gene-removed data respectively. Supplementary Figure S1 shows the P-network of 2616 genome isolates based solely on rRNA genes at *t* = 6. Although most phyla are connected to each other (i.e. 2613 connected nodes and z = 1321 at *t* = 6), we observed a clear separation between Archaea and Bacteria. These results suggest that rRNA genes can be used to infer a phylogeny that distinguishes Archaea from Bacteria, but these sequences do not provide sufficient resolution of various Bacteria phyla.

**Figure 2:**
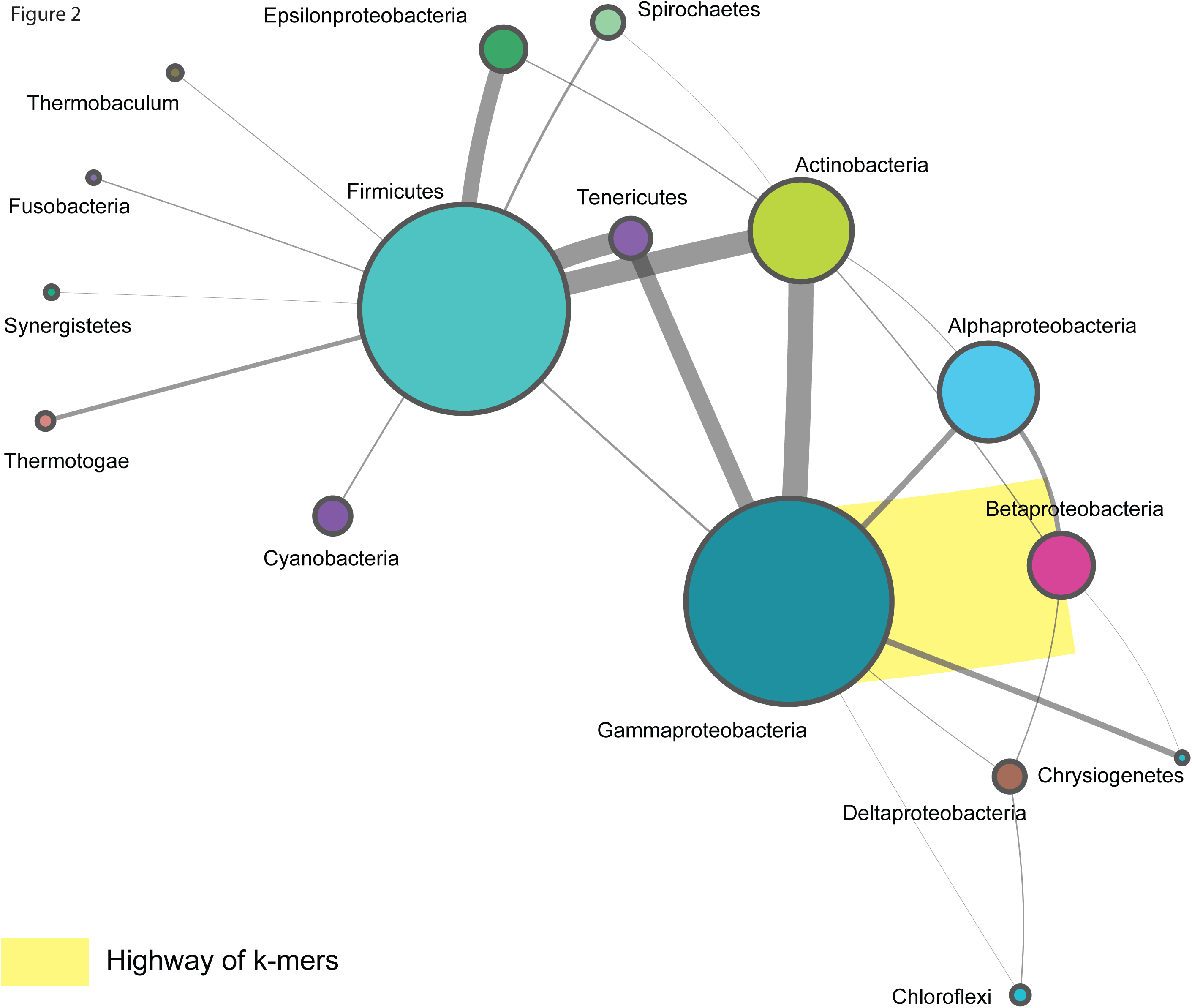
*P*-network of prokaryote genomes using 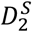 with *k*=25, based on whole-genome data with rRNA genes removed, at *t* = 3. An edge between two nodes represents the number of connections between isolates from the two phyla. The size of a node is proportional to the number of isolates within the phylum. Singletons are not shown.

### Evolution of plasmid genomes

To compare the evolutionary histories of extrachromosomal plasmids against those of whole genomes, we computed *I*- and P-networks using plasmid-only genome data for 921 isolates from 26 phyla (see Methods). Figure 3 shows the *I*-network of the 921 plasmid genomes at *t* = 0, in which *E* = 14.3 ( *c* = 745, *e* = 10679 and *z* = 48; Supplementary Table S5). Most phyla appear as distinct cliques, but notably with edges between the Actinobacteria, Firmicutes and the different classes of Proteobacteria. At *t* = 4 most phyla are separated as distinct cliques, with the exception of ε-Proteobacteria and Firmicutes; the other Proteobacteria (α, β, δ and γ) are in a distinct paraclique. The Euryarchaeota, connected only to the bacterial phylum of Planctomycetes at *t* = 0, is separated from Bacteria at *t* ≥ 1. All phyla are disjoint at *t* = 7. These results are not surprising, as the plasmid genomes are known to evolve faster than the core genomes, and in combination with their smaller genome size, fewer shared *k*-mers are observed at a high similarity threshold^26^.

**Figure 3:**
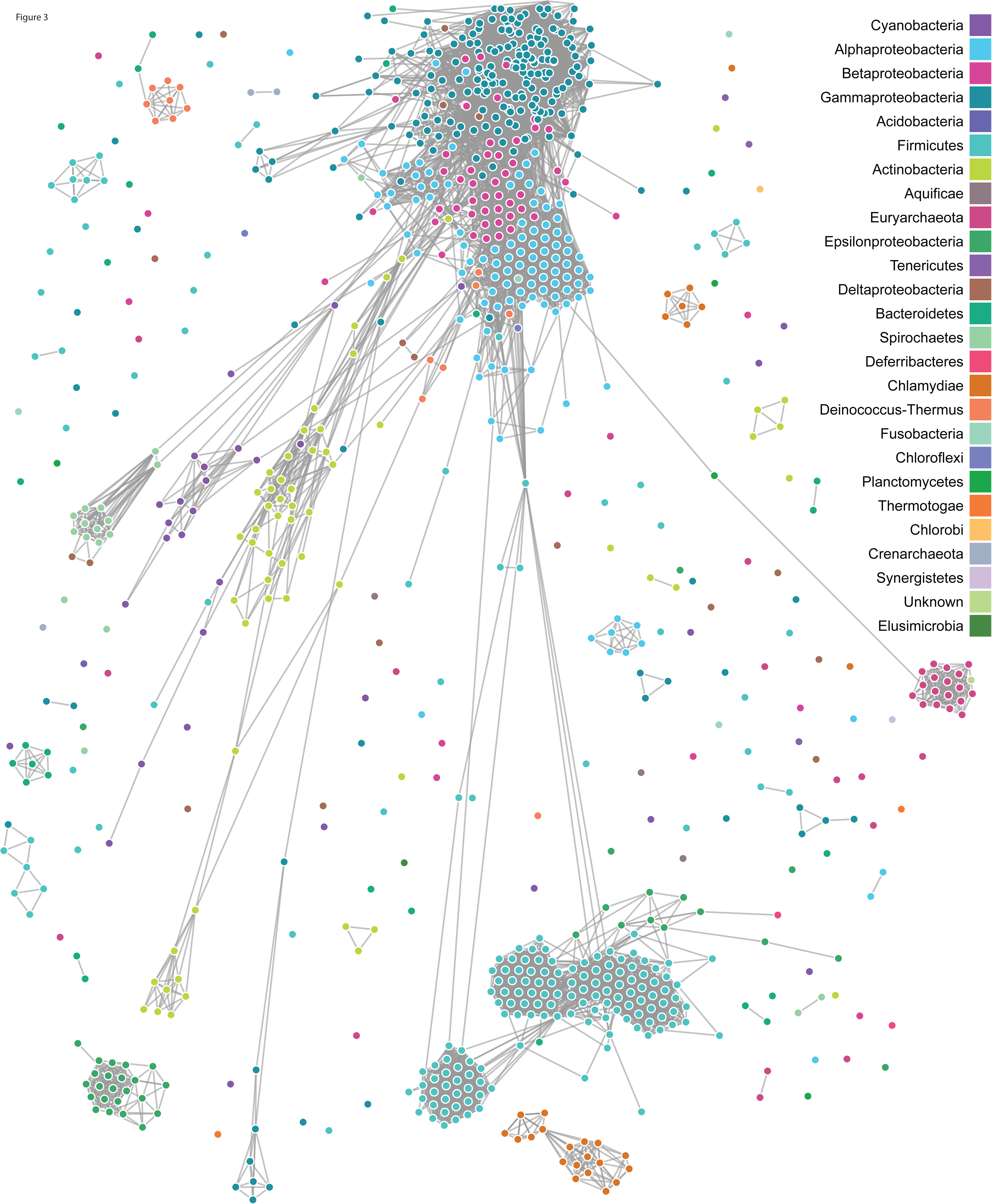
*I*-network of 921 plasmid genomes using 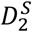 with *k*=25. An edge between two nodes represents evidence of share k-mers.

Figure 4 shows *E* for all four *I*-networks at different thresholds. For all networks, the number of edges per node (thus network density) decreases as *t* increases; a higher proportion of shared *k*-mers is required at a higher (more-stringent) threshold. The rRNA gene-only network is denser than the others, i.e. *E* remains >1000 for *t* = 0 through *t* = 6, compared to *E* < 200 in the others for *t* = 0 through *t* = 4. As expected, the highest density of the complete-genome network is observed at *t* < 2, *E* > 1400, and *E* decreases rapidly at *t* between 2 and 5. The network without rRNA genes has a lower density, *E* < 800, at *t* = 0 and decreases to level similar level to that of previous network at *t* = 5, e.g. *E* < 100. These results confirm that rRNAs are more highly conserved (i.e. the sequences are more similar as captured by 25-mers) than are the genome sequences overall. The density variation of the networks inferred based on whole-genomes and rRNA gene-removed data are more similar than the one observed for the rRNA-gene network. Figure 4 also shows that the plasmid network has the lowest density, *E* < 20 at *t* ≥ 0, implying that the plasmid genomes have diversified in 25-mer composition more rapidly than have the corresponding main genomes.

**Figure 4:**
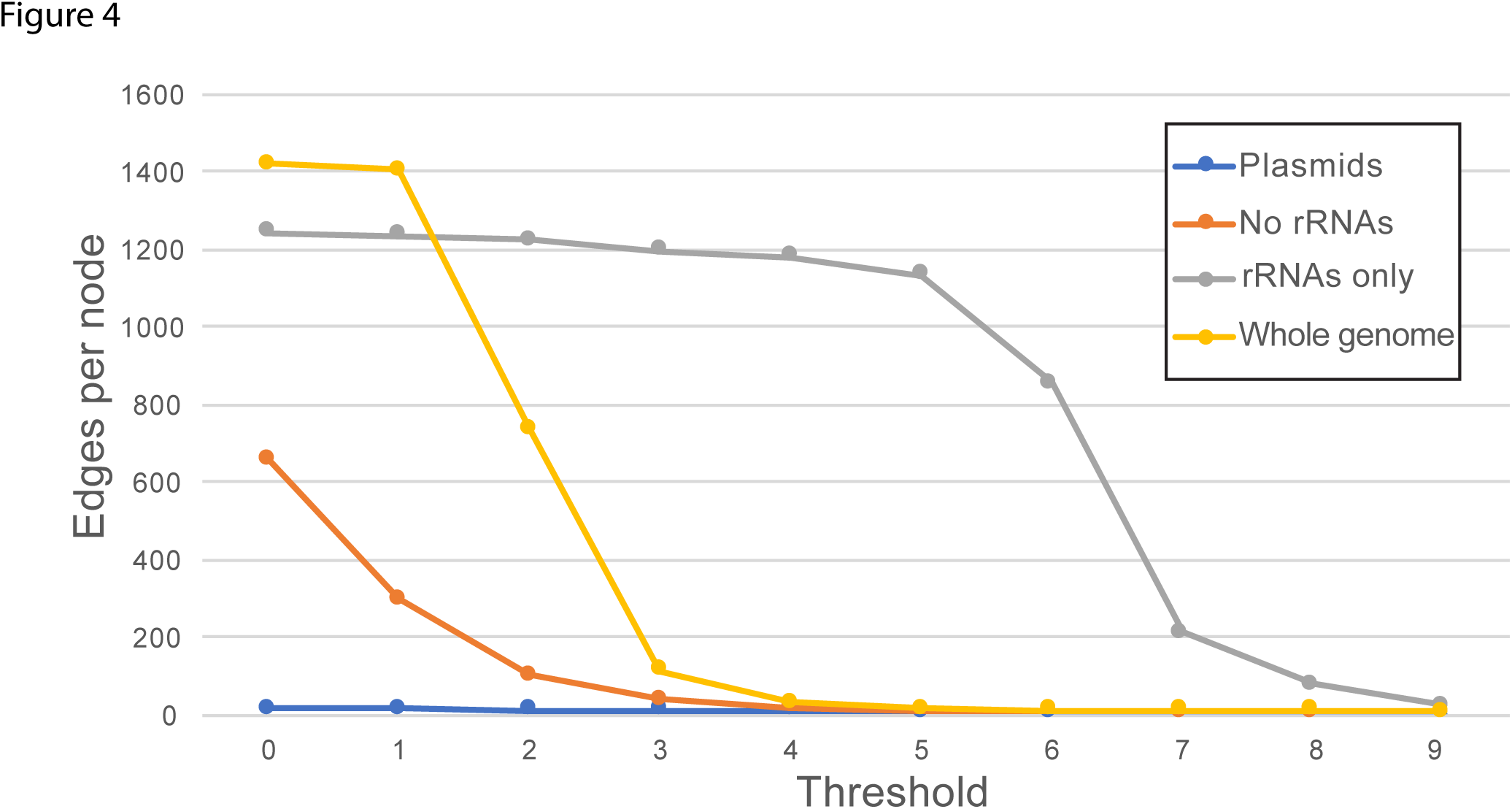
Number of edges per node, *E*, across distinct threshold levels of *t* for each *I*-network based on (a) complete genomes (core-genome with rRNAs + plasmids), (b) rRNA gene sequences, (c) complete genomes without rRNA genes, and (d) plasmid genomes.

### Core *k*-mers of microbial genera

We define a *core k-mer* in a group of interests as a *k*-mer that is present in every genome within the group, *e.g*. a core 25-mer in Proteobacteria is present in all proteobacterial genomes in our database (see Methods). Here, we identified core 25-mers for each genus in our 2785-genome dataset. Of these 699 genera, 497 consist of only a single genome isolate, and 51 consist of highly divergent genomes for which no core 25-mers were identified; we exclude these data from this part of analysis. The 151 genera for which core 25-mers were identified are shown in Supplementary Table S6. To represent the variable numbers of representative isolates of these genera in our dataset, we define *K* as the number of distinct core *k*-mers per isolate for each genus; this value can indicate the extent of genome divergence (and thus evolutionary rate of these genomes) for each of these genera. The three genomes of *Azotobacter* have the highest number of core *k*-mers, with *K* = 1722079; these genomes represent distinct isolates of the same species, *Azotobacter vinelandii*. This is in contrast to the 123 *Streptococcus* genomes (of 27 species) that share only one core k-mer (K = 0.008). Among the 20 genera with the greatest *K* values, *Shigella* has the highest number of distinct isolates (10 isolates from four species) at K= 33698. This is in stark comparison to *K* = 4.82 among the 11 *Ralstonia* genome isolates from three species. Thus these *Shigella* genomes have diverged less from their common ancestor than have these *Ralstonia* genomes from theirs, as assessed by shared 25-mers.

### Core functions of microbial phyla

To relate the shared *k*-mers to biological functions, we assembled all 25-mers in the 2785 genomes and their associated genome locations and annotated function based on Clusters of Orthologous Groups (COGs^27^) in a relational database. Then using the core 25-mers above, we identified the core functions in each of the 151 genera based on annotated functions that are associated with these *k*-mers (e.g. using k-mer position information to identify the corresponding gene, or non-coding sequence, and the gene annotation when available). For this analysis, we focused on protein-coding sequences (i.e. rRNA sequences were discarded: Methods), resulting a set of core 25-mers from 112 genera in 15 phyla; the corresponding COG functional categories for these core 25-mers are shown in Supplementary Table S7. The non-informative functional categories R (General function prediction only) and S (Function unknown) were excluded in subsequent analysis. We do not identify any core *k*-mers related to the functional category Y (Nuclear structure) in our dataset. The less-represented functional categories in our data (those with proportion <1%) are A (RNA processing and modification), B (Chromatin structure and dynamics), W (Extracellular structure) and Z (Cytoskeleton). The Chloroflexi, Euryarchaeota and Thaumarchaeota are the only phyla with evidence of core *k*-mers associated with the functional category B. The functional category A is related to core *k*-mers only in the proteobacterial classes (with the exception of the ε-Proteobacteria) and in phylum Actinobacteria. Figure 5 shows the proportions of COG number associated with core 25-mers across the 23 COG categories for 16 phyla, showing the top five categories for each phylum. Categories E (Amino acid metabolism and transport) and C (Energy production and conversion) are among the top five categories in 15 and 13 phyla respectively. The ε-Proteobacteria, Thaumarchaeota, Euryarchaeota, Actinobacteria, Cyanobacteria and Chloroflexi are also the only phyla with category H (Coenzyme metabolism) in the top five. For the phyla Tenericutes, Deinococcus-Thermus, Firmicutes and Crenarchaeota, the most represented functional categories include P (Inorganic ion transport and metabolism), L (Replication and repair), J (Translation), E and G (Carbohydrate metabolism and transport). The phylum Bacteroidetes is the only phylum for which categories O (Post-translational modification, protein turnover, chaperone functions), Q (Secondary structure) and F (Nucleotide metabolism and transport) are among the top five. Phylum Spirochaetes is the only one with U (Intracellular trafficking and secretion) and T (Signal transduction) in the top five, but the COG numbers associated with core 25-mers are extremely low.

**Figure 5:**
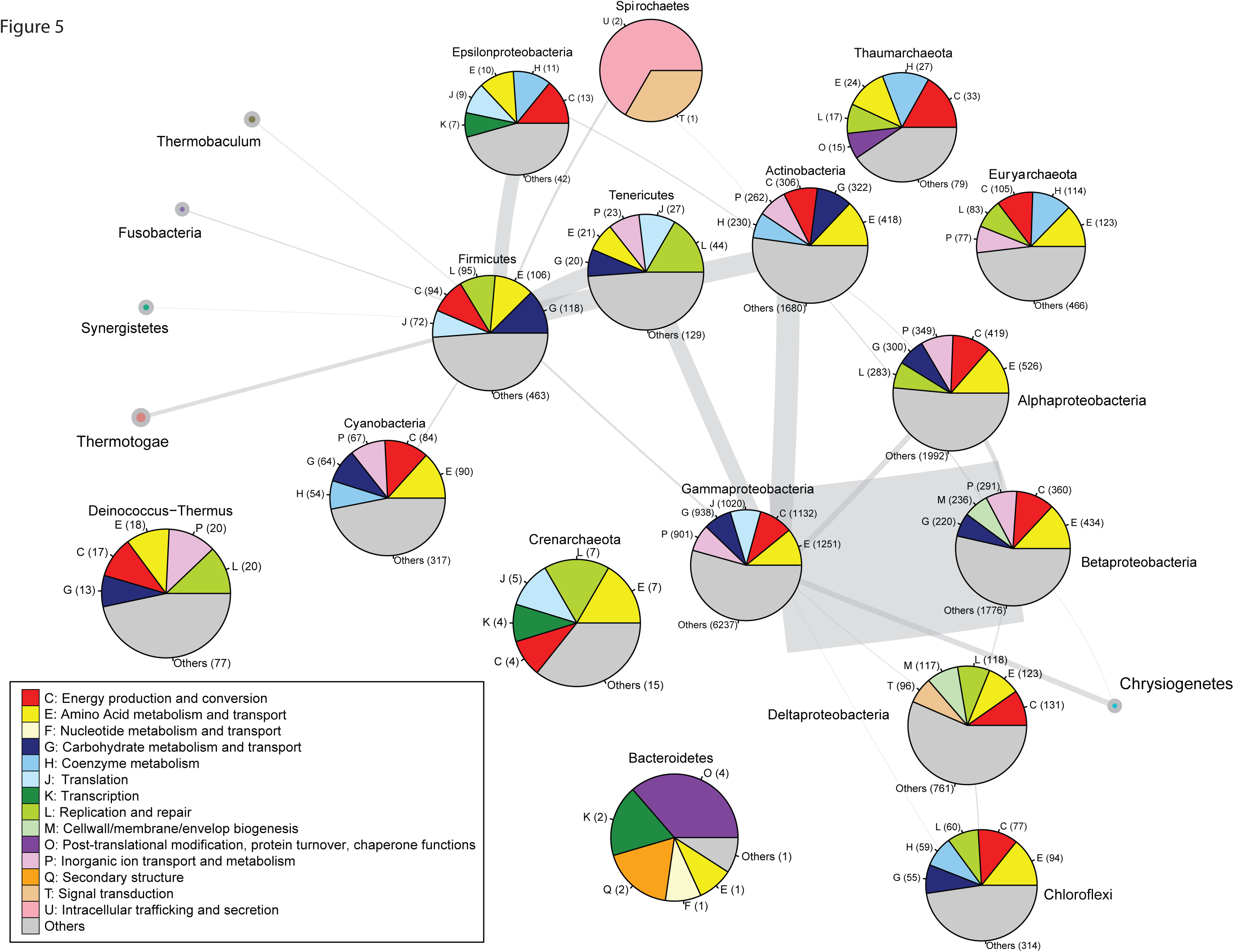
*P*-network of prokaryote genomes using 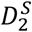 with *k*=25, based on whole-genome data with rRNA genes removed, at *t* = 3. The nodes are pie-charts representing the COG-category profiles of each phylum. Each COG category is color-coded. Only the top five categories are displayed; in most cases the top 5 categories account for at least 50%.

In order to find if the phyla can be clustered based on their COG categories profiles, we performed a series of PCA analysis. PCA on the raw data (e.g. non-normalised counts of COG number) did not show evidence of any particular clustering (Supplementary Figure S2). Nor do the genera cluster according to the number of isolates (Supplementary Figure S3). These results confirm that the different numbers of isolates per genus do not bias our analysis of functional categories. Supplementary Figure S4 shows the PCA analysis performed on the normalised counts of COG numbers with center-scaled COG categories (e.g. COG categories with equal weights). In this analysis *Nitrosopumilus*, the only genus in phylum Thaumarchaeota in this dataset, is isolated from the other genera. Genus *Dehalococcoides*, a member of phylum Chloroflexi, is likewise separated from the other genera by this measure.

### Computational efficiency and scalability

To compute the 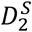 distance between microbial genomes we used a modified version of our own implementation of the *D*_2_ statistics^7^. This newer version was used to compute the 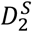 distance of two genomes at a time. Each pairwise distance can be computed independently, so we ran thousands of parallel jobs for each pairwise comparison for our different AF networks across a high-performance distributed-memory computing cluster. On average, it takes about five seconds to compute the 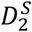 distance between two microbial genomes (time can vary depending on the genome size). The principal advantage of this approach is that it is not limited by memory consumption, as each job requires only a few hundred MB. Although visualisation of the network using the D3 library is scalable to large data, it can take a few minutes for the force-directed algorithm to provide an optimal layout for a densely connected network.

Extraction of the core *k*-mers took less than an hour for our dataset of 2785 microbial genomes. Mapping the core *k*-mers of 1475 genomes to our SQL database (store on a SSD hard-drive) took less than one hour.

## Discussion

In this study we demonstrate that AF approaches can be used to infer phylogenetic networks quickly and accurately for large-scale microbial whole-genome data. We introduce for the first time the concept of *k*-mer similarity network and two different types of AF networks, the *I*- and *P*-networks. We show that by combining a *k*-mer approach with the use of a relational database, biological information can be accessed for large-scale data at unprecedented speed. Finally, we define core *k-* mers as *k*-mers present in every isolate genome of a genus, following the concept of core genes^28^,^29^.

We examined the impact of rRNA genes and plasmids on the phylogenetic signal captured when computing phylogenomic relationships among microbial genomes. As expected, the rRNA genes contribute to the phylogenetic signal captured by 25-mers, as they do in MSA-based approaches. However, the pattern of network density versus threshold value (see Figure 4) indicates that the phylogenetic signal recovered here is not driven by rRNAs alone. Our result that, in general, these rRNA genes do not resolve relationships among (and sometimes within) bacterial phyla is in line with many previous studies^20,30-32^}. The density of the AF plasmid network confirms the large diversity of these mobile genetic elements, and we found similarity between the connections observed between this network and the ones based on whole-genomes, with or without rRNA genes, and rRNA gene sequences. The proteobacterial classes tend to have the strongest connections in all our AF networks, in particular between β- and γ-Proteobacteria, and we also observed strong similarity between the Actinobacteria and Proteobacteria or Firmicutes across all networks. The large extent of LGT between β- and γ-Proteobacteria^33^ isolates partly explains this strong similarity in our AF networks.

Overall, we demonstrated that the *I*- and *P*-networks provide a quick overview of the evolutionary relationships among whole genomes, or subsets of genomes, in large-scale datasets. Moreover, our AF networks, based on 25-mers pairwise comparison between two isolates, can be used to study the evolutionary dynamics aggregated at different taxonomic levels: by varying the distance threshold we can visualise evolutionary patterns among kingdoms (e.g. Archaea and Bacteria at *t* < 3), phyla (e.g. Proteobacteria, Firmicutes etc. at 3 ≤ *t* ≤ 5), classes (e.g. of Proteobacteria 4 ≤ *t* ≤ 6), and between and/or within genera (e.g. *Escherichia coli* and *Shigella* at *t* > 6).

Our approach to find the most highly conserved functions (apart from those of rRNAs) using core 25- mer profiles has shown that the biological functions associated with the metabolism and transport of amino acids, and the production and conversion of energy, are the categories most conserved in our dataset. The core 25-mer profiles revealed that similar core biological functions profiles are observed for phyla that share a large extent of *k*-mers in our AF network. Our analysis also indicates that the functions highly conserved in ε-Proteobacteria and in δ-Proteobacteria are distinct from those conserved in the other proteobacterial classes. Except for the two most highly conserved categories (above), the ε-Proteobacteria do not share highly conserved functions with the other classes of Proteobacteria; indeed, the ε-Proteobacteria share more 25-mers with the Firmicutes and with the Actinobacteria than with other Proteobacteria. These results support previous findings showing that the ε-Proteobacteria are the most-basal Proteobacteria by most criteria, and the last class among in this phylum to have been recognised^34^. Finally, we also observed that the Tenericutes are among the only phyla that do not have highly conserved functions related to energy production and conversion; this can be related to their parasitic lifestyle^35^.

Of these 699 genera, no core 25-mers were recovered for 51, particularly in genera represented by genome sequences for many isolates from different species. For these, a core *k*-mer sets might be sought at lower values of *k*, although at the potential risk of capturing a phylogenetic signal due to false positives and background noise. Similarly, some phyla that we used to identify highly conserved functions have few distinct COGs related to core 25-mers.

A major advantage of AF approaches in general (and this approach in particular) lies in its computational performance in the inference of phylogenetic networks, and the extraction and mapping of core *k*-mers to biological functions^7^,^36^. Because our approach consists of independent pairwise comparisons we can distribute the computation across multiple processors, greatly minimizing problems potentially arising due to demand on memory^7^. Here we inferred 25-mer similarity networks among < 2700 genomes in a matter of hours. To map core *k*-mers to our database we took advantage of the SQL architecture, indexing and hashing to compare billions of *k*-mers in a few minutes using an SSD hard drive. The database itself could be generated in only a few hours from RefSeq data of more than 4000 microbial isolates.

It would be of great interest to be able to discriminate the edges in the AF networks based on the dominant phylogenetic signal observed (e.g. vertical versus lateral). To visualise large phylogenetic network such as the one presented here, the D3 library (and web technology more generally) might not be the most optimal approach. Indeed, even with recent improvements of the JavaScript-based application and an optimised library such as D3, it is difficult for web browsers to display large networks in a force-directed layout. We could use instead use software specifically designed for visualisation of large networks, e.g. Gephi^37^, although undoubtedly at the expense of accessibility e.g. through unfamiliarity among users, or loss of cross-browser compatibility. Finally, we understand that an open-access, publicly available version of a *k*-mer database would be useful for our research community; however, such a database would require dedicated servers, management and support to be durable and useful at long-term for the community.

## Methods

### Data

The 2785 completely sequenced genomes of Bacteria and Archaea were downloaded from NCBI on 31 January 2016 (Supplementary Table S1); functional annotation of these genomes was obtained through the corresponding RefSeq records. Genes encoding ribosomal RNAs were identified based on annotation. Genomes with no annotation information were excluded from our rRNA genes network. Among the 2785 isolates, 921 contains plasmids; these plasmid genomes were used in the plasmid-only network.

### Relational database of *k*-mers and genome features

We extracted 10,059,526,408 distinct 25-mers from the genomes of 4401 bacterial and archaeal isolates (as of 31 of January 2016 in NCBI RefSeq), of which 2781 genomes are complete. We tabulated these *k*-mers, and their genomic locations and features (based on RefSeq annotations), in a relational database using SQL, following Greenfield and Roehm^36^. Tables in this database contain the list of isolates, the list of genes and their sequences, taxonomic information for each isolate, an indexed list of all 25-mers, an indexed list of gene-by-gene comparisons for each pair of genes, and an indexed list of genome-by-genome comparisons for each pair of genomes.

### AF network

We followed Bernard et al.^18^ in generating the AF networks. First we computed pairwise comparisons for the 2785 isolates and generated for each comparison the corresponding 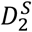 distance^7^ *d*, using 25-mers across parallel CPUs. For each pair of genomes *a* and *b*, we transformed *d* into a similarity measure *S_ab_*, where *S_ab_* = 10 – *d*. We discarded all instances for which *d* >10, as these pairs of sequences share ≤ 0.01% of 25-mers (i.e. there is almost no evidence of homology). We then generated the networks using JSON files containing the *S* values as input for a Javascript script using the D3 library (https://d3js.org/). Here, we present two types of AF networks. For a phylum-level depiction of the network ( *P*-network) we grouped all sequences of the same phylum as a single entity prior to calculating the distance; each phylum is represented by a node in the network. The width of the edge between two nodes represents the number of connections between isolates from these two phyla, and the size of each node is proportional to the number of isolates in the phylum. For an isolate-level depiction of the network (*I*-network) we treated each genome isolate as a single entity (i.e. node). In this network, an edge between two nodes indicates evidence of shared *k*-mers. The AF networks include a similarity-score threshold *t*, for which only edges with *S* > *t* are displayed; changing *t* therefore can dynamically change the structure of the network^18^. The resulting dynamic networks can be visualized using any web browser. All the networks can be found here: http://espace.library.uq.edu.au/view/UQ:543037^25^.

### Core *k*-mers and COG categories

For a specific group of microbial isolates (e.g. a genus, or a phylum) we extracted the set of 25-mers that are found in all isolates within the group; we define this set of 25-mers as the *core *k*-mers* for the corresponding group. Using the relational database of *k*-mers (above), for these core 25-mers we identified their corresponding genome locations and function based on COG (Clusters of Orthologous Groups)^38^ annotation in RefSeq records. We generated profiles of COG functional categories for each of the 151 genera, for each of the 11 phyla, and for the five proteobacterial classes in which core *k*-mers are identified using our approach.

### Computational scalability and runtime

Assessment of computational scalability was carried out using a high-performance distributed-memory computing cluster based on Intel Xeon Haswell (3.1 GHz) cores. Comparative runtime analysis of alignment-free methods was made on Intel Xeon Haswell E5-2667 v3 cores rated at 3.1GHz, using a single processor and one thread.

## Acknowledgements

We thank funding support from the Australian Research Council (DP150101875) awarded to MAR and CXC, and a James S. McDonnell Foundation grant awarded to MAR. This work was supported by computational resources of the National Computational Infrastructure (NCI) National Facility systems through the NCI Merit Allocation Scheme (Project d85).

## Author Contributions

G.B. implemented the analysis workflow and conducted the experiments, prepared all figures and tables, and the first draft of the manuscript. P. G. provided the *k*-mer database and contributed to analyses using this database. G.B., C.X.C. and M.A.R. conceived the study, designed the experiments, analysed and interpreted the results. All authors prepared, wrote, reviewed, commented on and approved the final manuscript.

